# Functional characterisation of gut microbiota and metabolism in Type 2 diabetes indicates that *Clostridiales* and *Enterococcus* could play a key role in the disease

**DOI:** 10.1101/836114

**Authors:** Marina Mora-Ortiz, Alain Oregioni, Sandrine P. Claus

**Author notes:** Joint first authors (Marina Mora-Ortiz, Alain Oregioni).

## Abstract

There is growing evidence indicating that gut microbiota contributes to the development of metabolic syndrome and Type 2 Diabetes (T2D). The most widely-used model for T2D research is the leptin deficient *db/db* mouse model. Yet, a characterisation of the gut microbial composition in this model in relationship with the metabolism is lacking. The objectives of this study were to identify metabolomics and microbial modulations associated with T2D in the *db/db* mouse model. The majority of microbial changes observed included an increase of Enterobacteriaceae and a decrease of Clostridiales in diabetics. The metabolomics interrogation of caecum indicated a lower proteolytic activity in diabetics, who also showed higher Short-Chain Fatty Acid (SCFA) levels. In the case of faeces, the model identified 9 metabolites, the main ones were acetate, butyrate and Branched Chain Amino Acids (BCAAs). Finally, liver was the organ with more metabolic links with gut-microbiota followed by the Gut-Brain Axis (GBA). In conclusion, the interaction between Clostridiales and Enterococcus with caecal metabolism could play a key role in the onset and development of diabetes. Further studies should investigate whether the role of these bacteria is causal or co-occurring.

## Introduction

Human microbiome accounts for circa 3.8 × 10^13^ microorganisms in the 70 kg ‘reference man’, including some unicellular eukaryotes, viruses, archae and bacteria (Sender *et al*., 2016). The intestine hosts the highest proportion of microorganisms, the largest fraction of this microbiome being located in the colon (circa 10^11^ microorganisms/mL) (Burrows *et al*., 2015, Suau *et al*., 1999, Savage, 1977, Andersson *et al*., 2008, Bäckhed *et al*., 2005, Berg, 1996, Sender *et al*., 2016). The main phyla are Bacteroides and Firmicutes. The relationship between these microorganisms and the host in healthy conditions is normally described as mutualistic, where both sides benefit from each other’s activity (Bäckhed *et al*., 2005, Seksik *et al*., 2003, Dunbar *et al*., 2002).

Some of these mutualistic interactions include the microbial fermentation of non-digestible polysaccharides which results in the production of Short-Chain Fatty Acids (SCFAs). As a consequence of this mutualistic relationship, the host receives carbon and the bacteria gain a protective anoxic environment and regular provision of carbon and nitrogen-rich food (Wong *et al*., 2006, Brune and Friedrich, 2000, Bäckhed *et al*., 2005).

Gut microbiota also play an important role in lipid absorption and homeostasis by regulating Bile Acids (BAs) metabolism, which promotes lipid absorption through the small intestinal barrier and acts as signalling molecules in the liver to regulate the Farnesoid X Receptor (FXR activity) and its downstream targets. An example of this includes the association between a reduction in bacteria from the genus *Lactobacillus* with alterations of the deconjugation of a BA (tauro-ß-muricholic acid) and downstream modulations of FXR signalling pathways, which was linked with reduced body weight gain (Cariou and Staels, 2007, Li *et al*., 2013). The FXR, activated by BA, plays an important role in the regulation of lipid and glucose metabolism, hepatic autophagy and BA metabolism and synthesis (Nie *et al*., 2015, Martin *et al*., 2019).

Bacteria can also deprive the host from available methyl donors (choline or betaine) transforming them into unavailable methyls in the form of TriMethylAmine (TMA). Hepatic Flavin-containing MonoOxygenases (FMOs) oxidize the TMAs into TriMethylAmine-*N*-Oxide (TMAO) which is further excreted in urine. In a larger scale, these methyl compounds are essential for epigenetic regulation and a number of other crucial biochemical pathways. Thus, gut microbiota impact lipid homeostasis by modulating hepatic metabolism through host access to methyl donors (de Castro *et al*., 2013, Koeth *et al*., 2013, Wang *et al*., 2011, Romano *et al*., 2017, Mora-Ortiz and Claus, 2017). Finally, the gut microbiome was also found associated with metabolic syndrome, particularly with obesity and diabetes (Karagiannides and Pothoulakis, 2007, Yang *et al*., 2019, Aydin *et al*., 2018).

Metabolic syndrome is characterised by the co-occurrence of various cardiovascular risk factors, such as central obesity, dyslipidaemia, hypertension and raised blood sugar promoted by insulin resistance (Huang, 2009). It ultimately leads to obesity, cardiovascular diseases and Type 2 Diabetes (T2D). This last disease is a complex metabolic disorder characterised by insulin resistance which currently affects 90% of people diagnosed with diabetes (https://www.who.int/diabetes/en/). High circulating blood sugar levels systematically disturb organs resulting in kidney failure, nerve damage, blindness and development of cardiovascular diseases (Tai *et al*., 2015, Amin *et al*., 2010, Anavekar *et al*., 2004, Trautner *et al*., 1997). The rapid spread of diabetes worldwide urges us to develop a better understanding of environmental factors contributing to the onset and development of the disease which requires validated models of T2D.

In the context of T2D, low microbiota diversity was previously linked with higher prevalence of insulin resistant phenotypes and low grade inflammation, associated with a reduction of microbial butyrate (Le Chatelier *et al*., 2013, Cotillard *et al*., 2013). A low level of inflammation of visceral adipose tissue has been correlated with insulin resistance and obesity, due to the link between transcripts implicated in inflammatory control and the accumulation of macrophage crown-like structures (Gesta *et al*., 2006, Apovian *et al*., 2008, Shimobayashi *et al*., 2018, Janochova *et al*., 2019). Some mouse studies have suggested however that the direction of change is the opposite, and it is insulin resistance which leads to inflammation (Shimobayashi *et al*., 2018). Regardless of the direction of change, this indicates a correlation between the innate immune system and insulin resistance mechanism (Aydin *et al*., 2018). In keeping with these findings, we recently observed a drastic metabolic modulation of the spleen in a mouse model of T2D (Mora-Ortiz *et al*., 2019). Despite growing evidence indicating that gut microbiota is a contributing factor to the onset and development of metabolic syndrome (a disorder originating from a dysregulation of energy metabolism and storage), the detailed links with the gut microbiota remain largely unknown.

One of the most widely-employed model for T2D investigation is the *db/db* (BKS.Cg-Dock7<m> +/+ Lepr<db>/ J) mouse model. Yet, only a few studies have looked at the gut microbiota of this model in association with metabolomics changes. Metabolomics’ references currently available do not include a characterization of caecal metabolism and faecal metabolomic changes over time (Saadat *et al*., 2012, Wei *et al*., 2015, Gipson *et al*., 2008, Connor *et al*., 2010, Mora-Ortiz *et al*., 2019). Our group previously characterised 18 relevant biological matrices of this mouse model, identifying over 60 metabolites associated with T2D (Mora-Ortiz *et al*., 2019). This study however, focussed on the metabolites from the host, and we did not cover the metabolites derived from the gut microbiota. Yet, these metabolites offer an invaluable window into the gut microbiota metabolic activity and provide an important insight into their role in the development of T2D. Here, we characterise for the first time the metabolic changes occurring in faeces over time and in the caecum content at 22 weeks of age, and their links with gut microbiota. Therefore, the objectives of this study were to i) evaluate gut microbiota metabolic changes in the *db/db* mouse model and compare stability of their metabolic activity over time, ii) characterise the gut microbiota at caecum level and iii) interrogate potential interactions between microorganisms from caecum and the host metabolism.

## Material and methods

### Animal handling and sample collection

Four-week-old mice (females, n=8; males, n=4) from the line BKS.Cg-Dock7<m> +/+ Lepr<db>/ J and their corresponding WT controls were bought from Charles River Laboratories, Italy. These animals were randomly allocated into two homogenous environments (according to their genotype, i.e. WT or *db/db*) and were acclimatised during one week upon arrival mixing bedding. After acclimatization, bedding was mixed on weekly basis according to the genetic background. Body weight was also measured at the same frequency. Faecal samples were collected at 7, 8, 9, 10, 12, 13, 14, 18 and 22-week-old (Figure 1.a). Caecum content was collected at end point during dissection, and immediately frozen into liquid nitrogen. Animals were euthanized by neck dislocation at week 22 in accordance with the regulations of the United Kingdom Animals (Scientific Procedures) Act, 1986 (ASPA) and approved by the Home Office (PPL 70/7942).

**Fig. 1:**
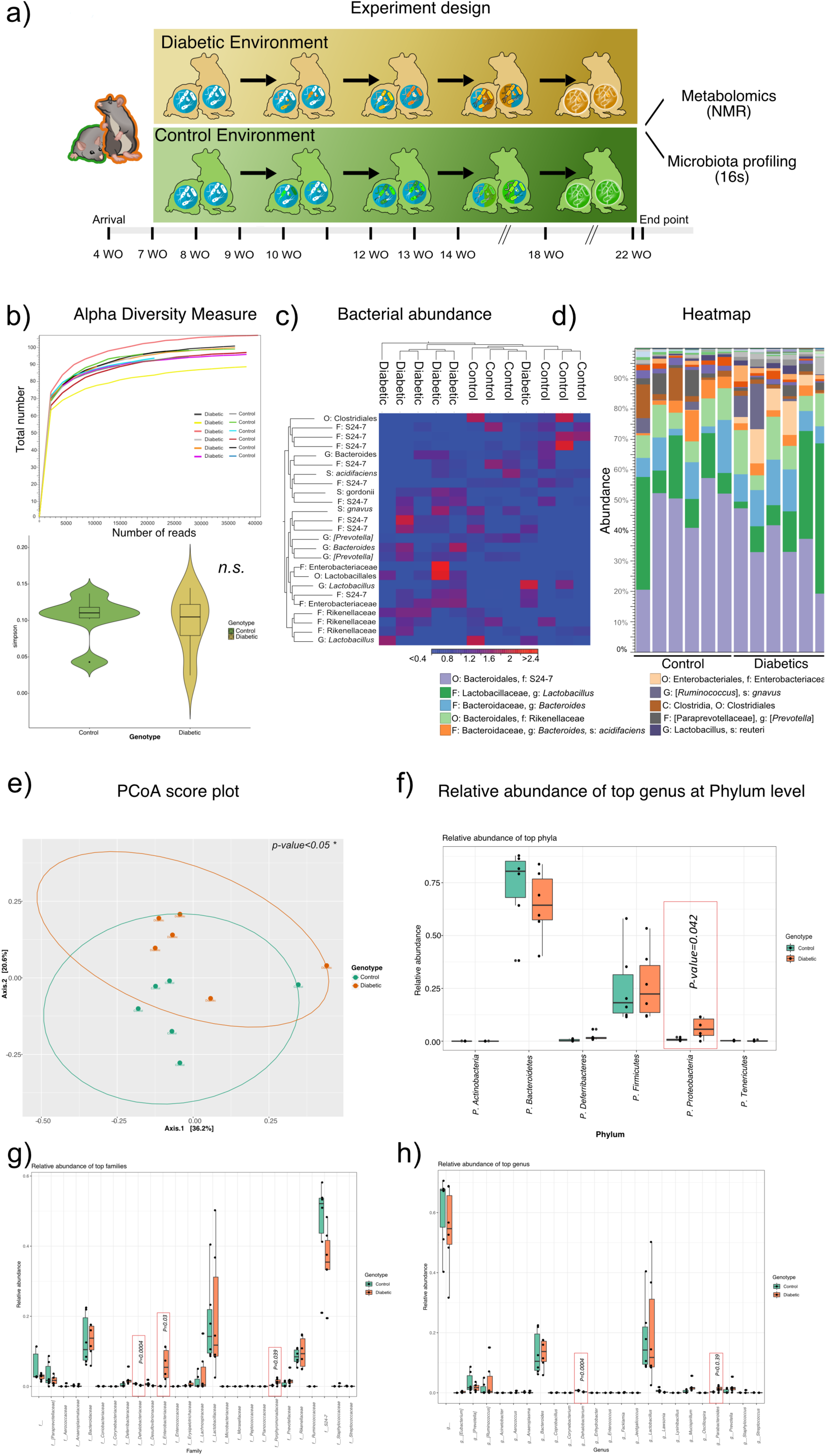
**a)** Twelve mice were allocated into two different homogenous environments, diabetic and control environment, according to their genetic background (*db/db*=6, with females=4 and males= 2; control=6, with females=4 and males= 2). **b) Top:** Alpha diversity rarefraction curve. **Bottom:** Alpha diversity violin boxplots grouped by genotype. **c)** Heatmap with clusters for diabetics (top left) and control (top right). **d)** Pile chart with OTU abundance results for control (left) and diabetic individuals (right). **e)** Beta diversity, groups were significantly different (*p*-value <0.05 *). **f)** The phylum Proteobacteria was significantly increased amongst diabetic. **g)** The family *Dehalobacteriaceae* was significantly reduced amongst diabetic individuals, while *Enterobacteriaceae* and *Porphyromonadaceae* were significantly increased. **h)** At genus level diabetic individuals had significantly lower abundance of *Dehalobacterium* and higher of *Parabacteroidetes*.

### DNA extractions, sequencing and analysis

DNA from faecal and caecum content samples was extracted using GeneMATRIX STOOL DNA Purification Kit (EURx Ltd, Poland). Aliquots of 10 μl with a concentration of 5 ng/μl were sent for 16s RNA paired end sequencing to the Welcome Genome Centre (University of Oxford, UK), where they were processed in an Illumina MiSeq System following the 16S Metagenomic Sequencing Library Preparation protocol. Briefly, the upstream 16s-pipeline was carried out in CLC Genomics Workbench (Qiagen Bioinformatics, Germany) and included i) merging of the paired reads, ii) fixed length trimming, iii) OTU clustering using Greengenes database and iv) alignment of the OTUs using MUSCLE. The downstream analysis was carried out in RStudio (version 1.2.1335) using the packages Phyloseq (McMurdie and Holmes, 2013) and Microbiome (Shetty *et al*., 2017, Leo Lahti, 2017) as well as MicrobiomeAnalyst (https://www.microbiomeanalyst.ca). These analyses included: i) alpha diversity analysis (interrogated with *t*-test), ii) beta diversity analysis (interrogated with Permanova), iii) analysis of the bacterial abundance (pile charts and heat map) and iv) study of the relative abundance of top taxa at phylum, family and genus level.

#### Caecum content and faecal waters

samples (∼30 mg) were homogenized utilising a TissueLyser LT (Qiagen, Germany) in 500 μl of phosphate buffer (0.2 M, pH 7.4)/D_2_0 (deuterium oxide (2,2-dimethyl-2-silapentane-5-sulfo- nate-d6 sodium salt)) (1:4) with 0.01%. The homogenates were centrifuged at 20,000 g during 15 min at 4 °C. A total of 200 μl of the supernatant was transferred into 3 mm NMR tubes for NMR analysis.

### NMR analysis

Bruker Avance HD 700 MHz NMR Spectrometer (Bruker Biopsin, Rheinstetten, Germany) equipped with a cooled SampleJet and a TCI CryoProbe from the same company was used to obtain all NMR spectra for faecal and caecum content waters. Standard 1D noesypr1d pulse sequences were used to acquire all one-dimensional (1D) NMR spectra, with a mixing time of 10 ms. A total of 64 scans were stored into 64 K data points; the spectral width was 13 ppm. Furthermore, 2D COSY experiments were carried out in all matrices.

### Metabolomics data pre-processing and multivariate statistical analysis

MestReNova version 11.0.2-18153 (Mestrelab Research S.L., Spain) was used to pre-process all spectra. Baseline corrections were carried out using Whittaker smoother algorithm and multipoint baseline correction when appropriate. Residual water (δ 4.70-5.10) and noise (regions before δ 0.5 and after δ 9.5) were removed. Calibration was done using TSP (δ 0.00) as reference. The spectra were then imported into Matlab version R2015b (Mathworks, UK) and analysed using Korrigan Toolbox version 0.1 (Korrigan Sciences Ltd., U.K.). Principal Component Analysis (PCA) was used to carry out a preliminary unsupervised study of the variability of the samples. This analysis was followed by a supervised pairwise Orthogonal Projection to Latent Structures Discriminant Analysis (O-PLS DA) which allowed the identification of modulations associated with T2D. Seven-fold cross-validation was used to evaluate goodness of prediction (*Q*^*2*^Y value) in the O-PLS DA model. O-PLS DA model were also used to investigate the correlation between the NMR spectra from 20 tissues or biofluids: caecum content, faecal content, plasma, spleen, kidney, liver, muscle, heart, hypothalamus, cerebrum, cerebellum, eye, duodenum, jejunum, ileum, proximal colon, transversal colon, distal colon, White Adipose Tissue or WAT apolar phase and WAT polar phase (previously published: (Mora-Ortiz *et al*., 2019)) and the bacteria abundance from the 16s analysis of caecum content and faeces respectively. Chenomx NMR Suite 8.2 from Chenomx Inc (Edmonton, Canada), online publically available databases such as the Human Metabolome Data Base (HMDB, http://www.hmdb.ca), the Biological Magnetic Resonance data bank (BMRB, http://www.bmrb.wisc.edu) and published literature were used to identify metabolites that were modulated.

### O-PLS DA model correlating NMR spectra from tissues and biofluids with bacterial abundance from caecum content

Bacterial abundance from caecum content (n=12) was correlated with 20 metabolomics’ matrices including: caecum content (n=12), faecal content (n=8), plasma (n=12), spleen (n=12), kidney (n=12), liver (n=6), muscle (n=12), heart (n=12), hypothalamus (n=12), cerebrum (n=12), cerebellum (n=12), eye (n=12), duodenum (n=12), jejunum (n=12), ileum (n=12), proximal colon (n=12), transversal colon (n=12), distal colon (n=12), WAT apolar (n=12) and polar phase (n=12) which were previously published (Mora-Ortiz *et al*., 2019). After a preliminary screening, only those models whose *Q*2 was positive and overfit was lower than 50% were considered for further analysis.

## Results and discussion

Previously, we published the metabolic and phenotypic profile of the same BKS.Cg-Dock7<m> +/+ Lepr<db>/ J and their corresponding WT control individuals (Mora-Ortiz *et al*., 2019). This study analysed 18 relevant biological matrices and identified over 60 metabolites in total. Here, we study the metabolism of faeces and caecum content, which was not previously described, and their relationship with gut microbiota, to get new insights into changes in gut microbes’ metabolic activity during the development of T2D.

### 16s rRNA sequencing analysis of caecum content at 22 weeks

The 16s rRNA sequencing analysis was carried out using *Illumina MiSeq System* targeting the V3 and V4 regions at 22 weeks of age, when diabetes is fully established. The rarefaction curve showed a total number of reads above 35,000 representing 85% of the total reads, which allowed an in-depth analysis of the microbiome (Figure 1.b, top part). The alpha diversity analysis per genotype did not show significant differences between groups (Figure 1.b, bottom part).

A heat map constructed from the OTUs abundance table showed clear clustering of diabetic individuals and controls. These clusters were principally driven by the presence of Clostridiales, *Bacteroides* sp. and Bacteroidales in controls (Figure 1.c). Diabetic individuals had lower relative concentrations of Clostridiales than controls, and higher levels of members from the family *Enterobacteriaceae* (Figure 1.d). Interestingly, *Lactobacillus reuteri* was four times higher in control individuals than in diabetics. Previous studies have shown that *L. reuteri* is associated with reduced levels of serum HbA1c and cholesterol in High-Fructose-Feed (HFD) rats, where it also had reduced insulin resistance (Hsieh *et al*., 2018, Hsieh *et al*., 2013).

Commensal clostridia, a group found reduced in diabetic individuals, plays an important role in gut homeostasis maintenance, producing butyrate that will be used by enterocytes promoting consequently gut health (Lopetuso *et al*., 2013, Pryde *et al*., 2002). Decrease in clostridia have also been previously associated with aging (Mikelsaar *et al*., 2010, Mäkivuokko *et al*., 2010, Zhao *et al*., 2011, Biagi *et al*., 2010). Thus, unbalanced levels of Clostridia and *Lactobacillus* could be linked with a dysfunctional gut microbiota among diabetic individuals, with potentially defective gut homeostasis and frailty.

Previous studies proved that altered microbiota in T2D patients was associated with reduction in butyrate-producing bacteria (such as *Clostridiales* sp., *Eubacterium rectale, F. prausnitzii* and *Roseburia* amongst others) and increases in *Lactobacillus* species such as *Lactobacillus gasseri (Qin et al., 2012, Karlsson et al., 2013)*, and showed a significant endotoxaemia, associated with the dysbiosis (Pussinen *et al*., 2011, Lassenius *et al*., 2011).

The beta diversity analysis showed significant differences between both groups (permanova *p*-value <0.05 *) (Figure 1.e). The analysis of the bacterial relative abundance indicated that the phylum Proteobacteria was significantly increased amongst diabetic. Previously, Larsen *et al*. (2005) showed that enrichment in members from the Proteobacteria phyla amongst diabetic individuals drived 28% of the PCA variability along with Actinobacteria (Larsen *et al*., 2010). Interestingly, LipoPolySaccharides (LPS) are some of the main compounds from the membranes of Proteobacteria and are inflammation promoters, which can produce endotoxaemia (Allcock *et al*., 2001), previously described as one of the main features associated with T2D.

In lower taxonomical levels, *Dehalobacteriaceae* was significantly reduced amongst diabetic individuals (*p*-value <0.001 **), while *Enterobacteriaceae* and *Porphyromonadaceae* were significantly increased (*p*-values <0.01 *). In a previous comparative study between *db/db* and *ob/ob* mouse gut microbiota, diabetic mice had *Dehalobacteriaceae* and *Enterobacteriaceae* enriched compared to the *ob/ob* model (Yang *et al*., 2019), which indicates different modulations inherent to each model and highlights the relevance to have model-specific microbial characterisations.

At genus level, the main changes occurred in the groups *Dehalobacterium* (*p*-value<0.001), which was decreased, and *Parabacteroidetes* (*p*-value<0.01), which was increased amongst diabetic individuals. Increases in *Parabacteroidetes* were also previously associated with high fat diet (Serino *et al*., 2012). Linear discriminant analysis Effect Size (LEfSe) results were consistent with the results described above. Further details are provided in S1.

### Metabolomics’ NMR analysis of caecum and faeces

Faecal pellets were collected at regular intervals to observe time-related changes concomitant with disease development. The O-PLS DA model using **faecal waters** as independent variables, and time points and genetic background as dependant variables (*R*^2^Ycum = 0.82 and a *Q*^2^Ycum = 0.34) showed quasi parallel trajectories, indicating that irrespective of the weekly homogenisation of soil litters, the metabolic shift associated with differences in genetic background remained constant. Diabetic individuals had higher levels of several amino acids (aspartate, threonine, leucine, BCAAs, histidine, phenylalanine and tyrosine), and lower levels of propionate and acetate, indicating that the faecal microbiome of diabetic individuals had a higher proteolytic and a lower saccharolytic activity (Figure 2, left panels). Increased levels of SCFAs were previously described associated to microbial fermentation of fibre in human and animal studies, showing an inverse relationship with body weight and adiposity (Lin *et al*., 2016, Chambers *et al*., 2018). Interestingly, higher levels of acetate in distal colon, which contributes to the regulation of central appetite (Frost *et al*., 2014), are considered more effective than in proximal colon. These have been shown to stimulate fat oxidation and improve inflammatory status and glucose homeostasis in men (van der Beek *et al*., 2016, Canfora and Blaak, 2017). A systematic decrease of acetate in faeces (more representative of distal colon) of diabetic mice could be linked with dysbiosis and with diabetic symptomatology.

**Figure 2.**
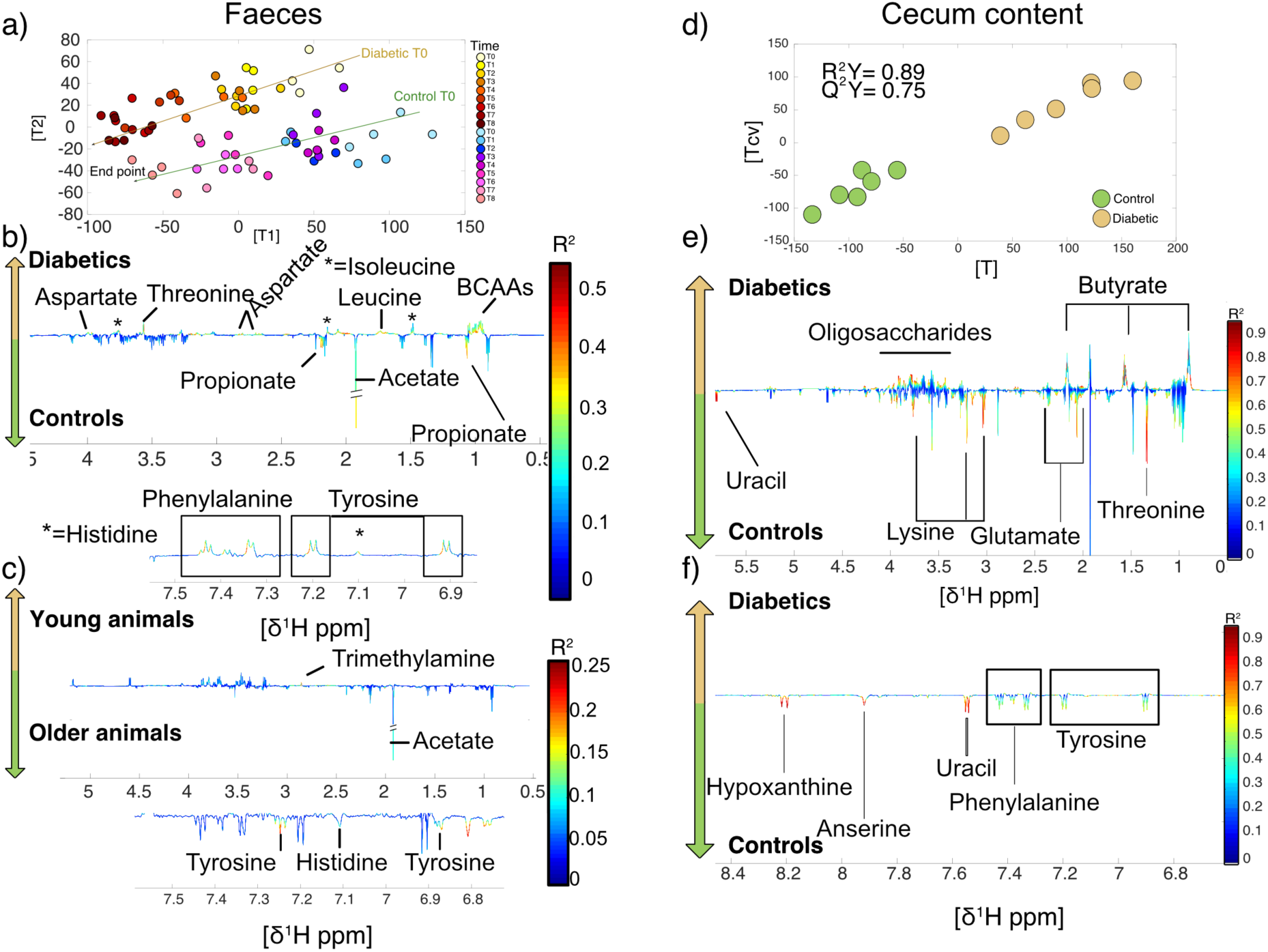
<u>Left hand panels:</u> faecal waters. **a)** Plot of the scores against the cross validated scores generated from the O-PLS DA model calculated using all spectra as a matrix (n=8) of independent variables and genetic background and time as predictors. **b & d)** Associated loading plots for genotype and age respectively. <u>Right hand side panels</u>: caecum content. **d)** Plot of the scores against the cross validated scores generated from the O-PLS DA model calculated using all spectra as a matrix (n=12) of independent variables and genetic background as predictor. **e & f)** Associated loading plots.

In the case of **caecum content**, the O-PLS DA model (*R*^2^Y= 0.89, *Q*^2^Y=0.75, n=12) showed that diabetic individuals had higher levels of butyrate potentially resulting from protein and oligosaccharides fermentation (Neis *et al*., 2015), and lower levels of uracil, lysine, glutamate, threonine, hypoxanthine, anserine, uracil, phenylalanine and tyrosine (Figure 2, right panels). SCFAs such as butyrate and acetate, are a source of energy for the enterocytes, and are linked to bacterial production (Pryde *et al*., 2002, Lopetuso *et al*., 2013), indicating higher levels of fermentation among diabetic individuals. Conversely, proteolysis seems lower among diabetic individuals, with lower levels of threonine and lysine. Proteolysis in the large intestine has been associated with *Clostridium, Bacteroides, Propionibacterium, Fusobacterium, Streptococcus* and *Lactobacillus*, being *Clostridium* especially relevant in T2D (Macfarlane *et al*., 1988, Qin *et al*., 2012).

The O-PLS DA model (*R*^2^Y= 0.99, *Q*^2^Y=0.95, n=10) comparing the metabolism between faeces and caecum of healthy individuals only (Figure 3) showed higher levels of SCFAs in faeces with higher levels of butyrate and acetate, and higher proteolytic activity in caecum, where levels of lysine, glycine, taurine, glycerol, glutamate, threonine, BCAAs and phenylalanine and tyrosine were higher. The higher levels of microbial metabolites such as phenylalanine and tyrosine in caecum was expected, since this organ contains a higher bacterial population (Dodd *et al*., 2017, Fujisaka *et al*., 2018). Interestingly, compared to humans, proteolysis is higher in the proximal colon than in the distal colon although the bacterial concentration is lower in the proximal colon. Previous studying comparing mouse and human models already pointed out significant differences between the intestinal system of mice and human (Hugenholtz and de Vos, 2018). Some of the most remarkable differences considered the pH range (human=glandular stomach with pH 1, mice= non-glandular forestomach with pH 3-4), the relative size of the small intestine (human=10 cm/kg, mice=1,500 cm/kg), and differences in the large intestine including the relative size and shape (Ghoshal and Bal, 1989, Hugenholtz and de Vos, 2018, Benson *et al*., 2010, Scholtens *et al*., 2012, Treuting and Dintzis, 2012). The microbiota of both species is similar at phylum level, being dominated by Bacteroidetes and Firmicutes, although we can also find differences here. For example, the phylum Deferribacteres is common in the mouse intestinal system and found very rarely in humans (Ley *et al*., 2006, Rawls *et al*., 2006, Bik *et al*., 2006). Studies comparing humans and 3 different mouse strains concluded that despite similarities in the profile between species, there are quantitative differences, despite many bacteria are shared between human and mice (approximately 42% of the core microbiome), only 4% of the genes are common in both species (Krych *et al*., 2013, Xiao *et al*., 2015). These discrepancies in microbiome could be partially responsible for the differences found in proteolysis, which is a feature to consider when interpreting the microbial results from mouse studies.

**Figure 3.**
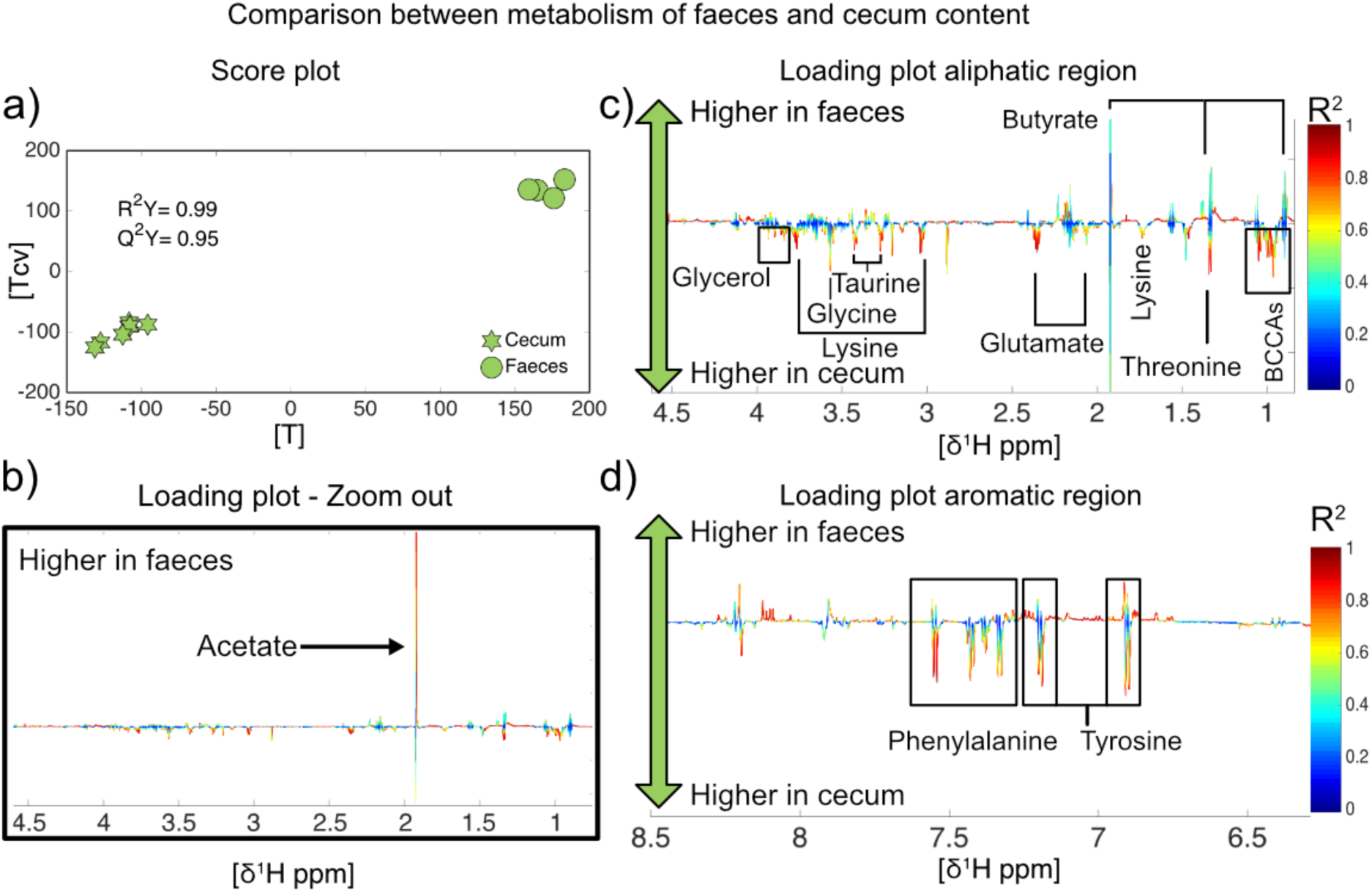
**a)** Plot of the scores against the cross validated scores generated from the O-PLS DA model calculated using all spectra as a matrix (n=10) of independent variables and location as predictor. **b & c)** Associated loading plot, aliphatic region of the ^1^H NMR spectra. **d)** Associated loading plot, aromatic region of the spectra.

### O-PLS DA correlation between metabolism and bacterial abundance in caecal content

Caecum bacterial activity was correlated with metabolomics changes from other tissues of the same individuals previously published (Mora-Ortiz *et al*., 2019). The O-PLS DA models were built using bacterial abundance from caecum as predictors and the metabolomics modulations as matrix of independent values.

In average, 36.8 bacteria per tissue and/or biofluid were found modulated and potentially involved in diabetes, with a standard deviation of 13.6 units. The tissues with lower number of valid O-PLS DA model was ileum, this could be due to a lower number of metabolic modulations in this tissue associated with diabetes (S2). Brain was the tissue with higher number of valid O-PLS DA models, with 62 models in total, however, it was overcome by liver when the *R*^2^ threshold was set up to 0.75. This could indicate that liver, possibly through its BAs interactions with gut microbiota is one of the main organs affected by the shift in microbiome (Figure 4 and S3). Interestingly, liver connexion with gut microbiota was mainly dominated by interactions with Bacteroidales, while brain and hypothalamus mainly linked mostly with Clostridiales (see figure 4 and S3).

Gut-Brain Axis (GBA) is also an area of particular interest in T2D research. GBA investigations have mainly covered diseases such as hepatic encephalopathy, depression, autism, functional gastrointestinal disorders and irritable bowel disease (Morgan, 1991, Foster and McVey Neufeld, 2013, Naseribafrouei *et al*., 2014, Mayer *et al*., 2014, Koloski *et al*., 2012). However, limiting literature is currently available describing GBA interactions in the context of T2D (Bessac *et al*., 2018).

Some of the most relevant bacteria involved in metabolomics interactions with tissues were *Lactobacillus*, Bacteroidales S24-7, *Parabacteroides gordonii, Mucispirillum schaedleri*, and Enterobacteriaceae, which had valid O-PLS DA models for metabolic interactions with spleen, kidney, liver, muscle and heart. The last two bacterial taxa were also associated with metabolic interactions in plasma. In the case of gut content (faeces and caecum content), only *Erysipelotrichaceae* and Bacteroidales S24-7 were common to both matrices.

*Erysipelotrichaceae* was again relevant in the GBA, and also Bacteroidales S24-7, Bacteroidales and *Rikenellaceae*. None of the bacteria was relevant across all the tissue sections of the gut, but *Anaeroplasma*, previously associated with diabetes (Brown *et al*., 2016), has the most promising perspectives with the highest number of valid O-PLS DA models. In the case of WAT, Bacteroides and Bacteroidales S24-7, Clostridiales and Mollicutes produced valid O-PLS DA models.

In total, 18 bacteria were found associated with metabolic perturbations in clusters of tissues or biofluids (note, eye was not considered in this summary, for more information about eye metabolic modulations linked to bacteria abundance see S2). More details about the models are provided in the heat maps of the supplements S4 and S5.

**Figure 3.**
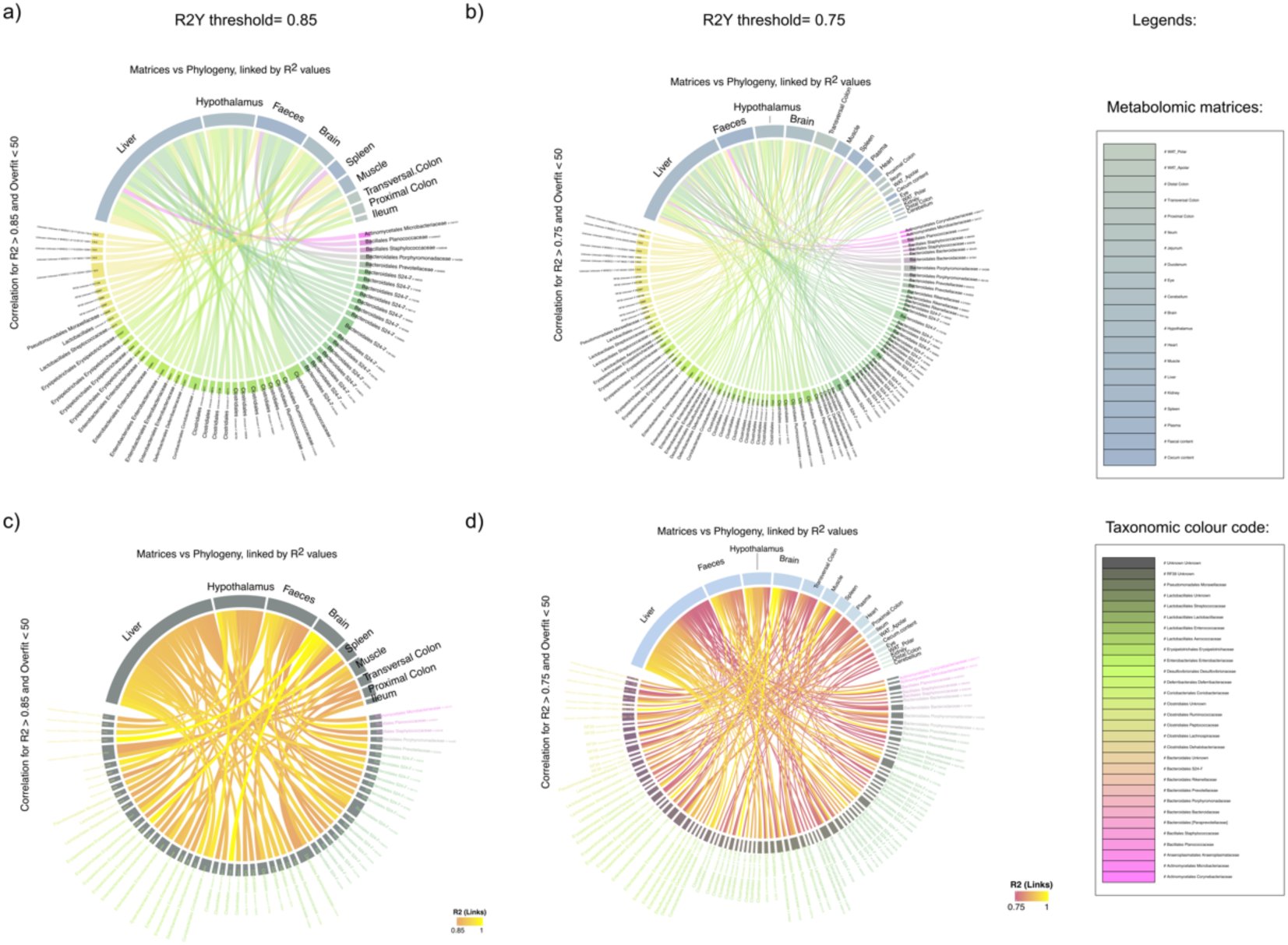
Chord diagram of the O-PLS DA models obtained comparing NMR matrices of independent values with bacterial abundance for each taxon identified as predictor. Only models with overfit smaller than 50% are considered. a & b) Taxa is colour coded. c & d) R2 values are colour coded. First column: R2Y threshold is set up at 0.85; second column: R2Y threshold is set up at 0.75. The Chord diagrams were plotted with the Circlize package under R (Gu *et al*., 2014).

## Conclusions

Differences in the abundance of Clostridiales and Enterococcus in caecum could indicate a link with the onset and development of diabetes. Further studies in T2D should interrogate whether differences in abundance between Clostridiales and Enterococcus are a cause or a consequence. For this purpose, germ-free mouse models, *in-vitro* or longitudinal human studies could be employed. Lower proteolytic activity was identified in caecum of diabetic individuals who had higher SCFAs, which are a result of the fermentation. Moreover, caecum was found to have more proteolytic activity among healthy individuals than lower sections of the colon; compared to humans, proteolysis was higher in the proximal colon. Finally, differences in gut microbiota associated with T2D mostly influenced liver and the GBA. Further studies should investigate if gut microbiota plays a causal role in this association.

## Abbreviations

BA: Bile Acids
BCAA: Branched-Chain Amino Acids
*db/db* mice: mice: BKS.Cg-Dock7<m> +/+ Lepr<db>/ J mice
NMR: Nuclear Magnetic Resonance;
TMA: trimethylamine;
TMAO: trimethylamine-*N*-oxide;
T2D: Type Two Diabetes;
O-PLS: Orthogonal Projections to Latent Structures;
SCFA: Short-Chain Fatty Acids.

## Acknowledgements

The authors would like to thank the Medical Research Council (MRC) for supporting this research (M004945/1). We also want to thank all the members from the Biological Resource Unit (BRU) from the University of Reading for their technical support, specially to Wayne Knight and Sophie Reid.

This work was also supported by the Francis Crick Institute through provision of access to the MRC Biomedical NMR Centre. The Francis Crick Institute receives its core funding from Cancer Research UK (FC001029), the UK Medical Research Council (FC001029), and the Wellcome Trust (FC001029).

## Availability of data and material

All NMR and 16s data are available in S2.

## Authors’ contributions

MMO conceived, designed and performed the experiments, analysed the data and wrote the manuscript; AO, led the NMR experiments and supervised the work, contributed to the analysis, the figures and the manuscript; SPC conceived and supervised the work and the manuscript. All authors read and approved the final manuscript.

