## Supplementary material for "Functional characterisation of gut microbiota and metabolism in Type 2 diabetes indicates that *Clostridiales* and *Enterococcus* could play a key role in the disease": S1

**Linear Discriminant Analysis (LDA) Effect Size (LEfSe)**

**Figure 1.** LEfSe results at phylum level:

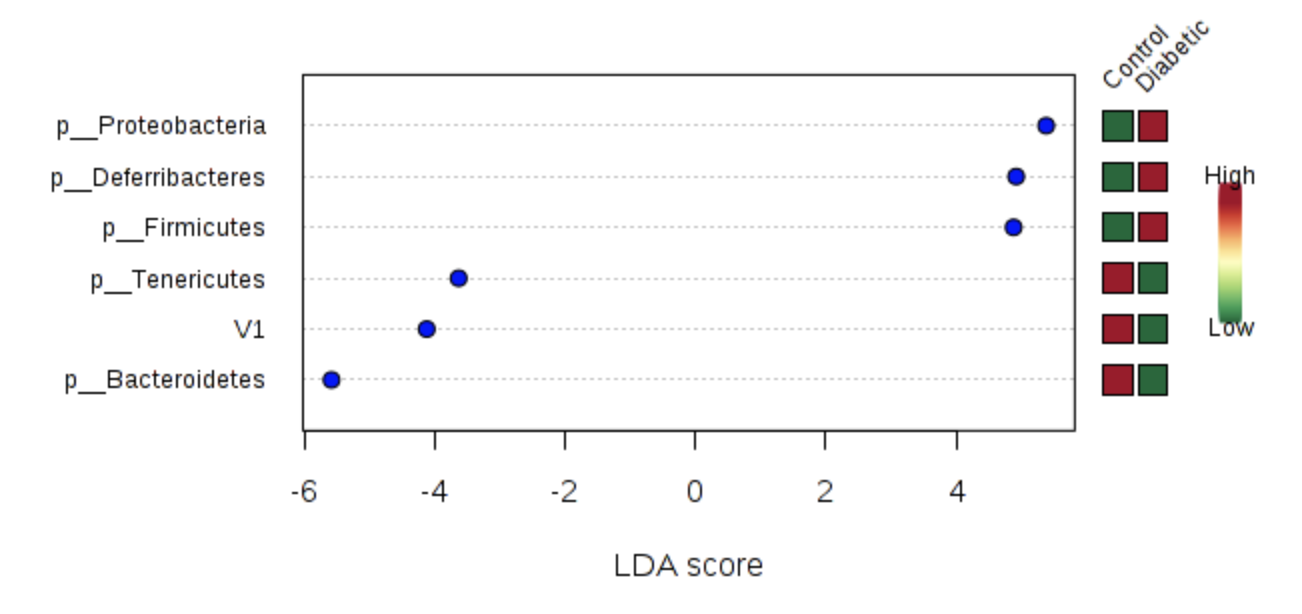

**Table 1.** LEfSe results at phylum Level:

|  | **Pvalues** | **FDR** | **Control** | **Diabetic** | **LDAscore** |
| --- | --- | --- | --- | --- | --- |
| **p__Deferribacteres** | 0.016309 | 0.097855 | 42654 | 210820 | 4.92 |
| **p__Proteobacteria** | 0.054664 | 0.16399 | 76443 | 551310 | 5.38 |
| **p__Tenericutes** | 0.20018 | 0.3935 | 24554 | 16097 | -3.63 |
| **p__Bacteroidetes** | 0.26233 | 0.3935 | 7196100 | 6436000 | -5.58 |
| **V1** | 0.74877 | 0.87278 | 150270 | 123990 | -4.12 |
| **p__Firmicutes** | 0.87278 | 0.87278 | 2510000 | 2661800 | 4.88 |

**Figure 2.** LEfSe results at family level:

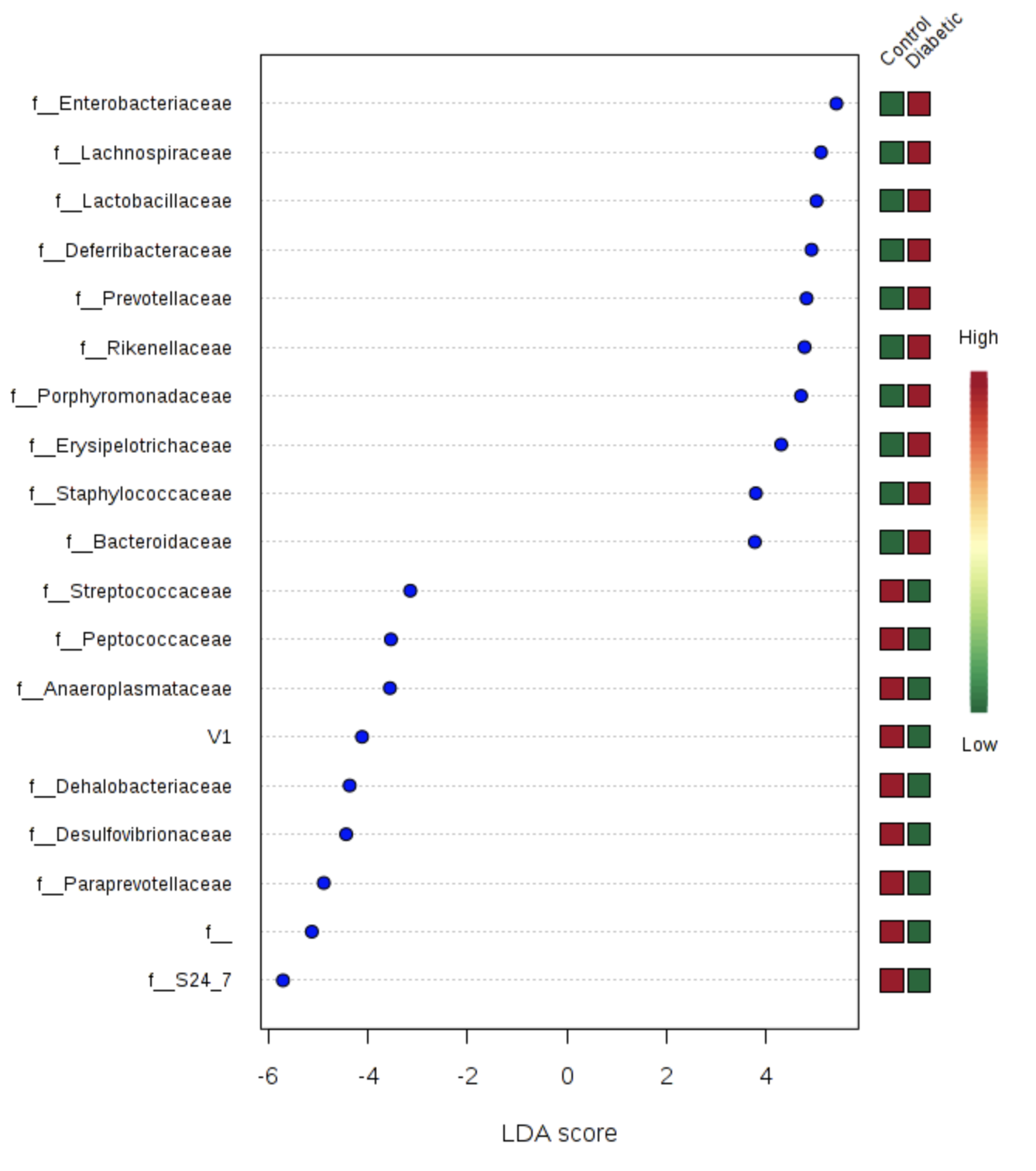

**Table 2.** LEfSe results at family level:

|  | **Pvalues** | **FDR** | **Control** | **Diabetic** | **LDAscore** |
| --- | --- | --- | --- | --- | --- |
| **f__Dehalobacteriaceae** | 0.0039478 | 0.075007 | 77270 | 30384 | -4.37 |
| **f__Deferribacteraceae** | 0.016309 | 0.11863 | 42654 | 210820 | 4.92 |
| **f__Enterobacteriaceae** | 0.020096 | 0.11863 | 2481.6 | 532750 | 5.42 |
| **f__Desulfovibrionaceae** | 0.024975 | 0.11863 | 73961 | 18559 | -4.44 |
| **f__Prevotellaceae** | 0.052976 | 0.1731 | 51717 | 184560 | 4.82 |
| **f__Porphyromonadaceae** | 0.054664 | 0.1731 | 27820 | 129270 | 4.71 |
| **f__S24_7** | 0.078169 | 0.21217 | 4582500 | 3560500 | -5.71 |
| **f__Peptococcaceae** | 0.10931 | 0.25962 | 10063 | 3159.8 | -3.54 |
| **f__Erysipelotrichaceae** | 0.14954 | 0.3157 | 14517 | 55145 | 4.31 |
| **f__Anaeroplasmataceae** | 0.22151 | 0.42087 | 21588 | 14402 | -3.56 |
| **f__Staphylococcaceae** | 0.2615 | 0.45168 | 3160.1 | 15863 | 3.8 |
| **f__Lachnospiraceae** | 0.50659 | 0.8021 | 130710 | 389540 | 5.11 |
| **f__Rikenellaceae** | 0.63095 | 0.83687 | 836690 | 956890 | 4.78 |
| **V1** | 0.74877 | 0.83687 | 150270 | 123990 | -4.12 |
| **f__Paraprevotellaceae** | 0.74877 | 0.83687 | 329380 | 175660 | -4.89 |
| **f__Lactobacillaceae** | 0.74877 | 0.83687 | 1812500 | 2023900 | 5.02 |
| **f__** | 0.74877 | 0.83687 | 535500 | 267970 | -5.13 |
| **f__Streptococcaceae** | 0.93607 | 0.98808 | 5503.3 | 2705.6 | -3.15 |
| **f__Bacteroidaceae** | 1 | 1 | 1291800 | 1303900 | 3.78 |

**Figure 3**. LEfSe results at genus level:

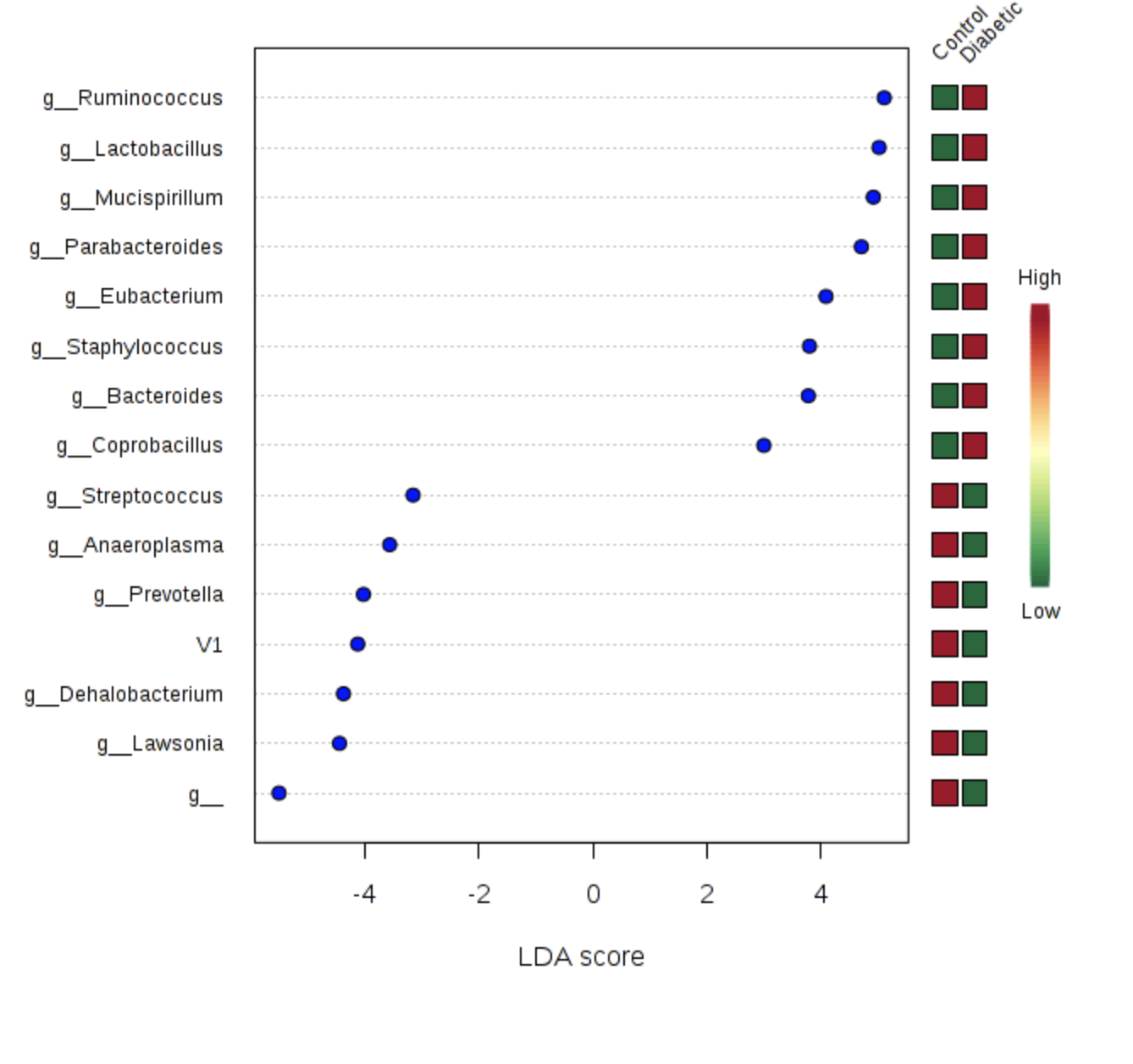

**Table 3.** LEfSe results at genus level:

|  | **Pvalues** | **FDR** | **Control** | **Diabetic** | **LDAscore** |
| --- | --- | --- | --- | --- | --- |
| **g__Dehalobacterium** | 0.0039478 | 0.059216 | 77270 | 30384 | -4.37 |
| **g__Mucispirillum** | 0.016309 | 0.12232 | 42654 | 210820 | 4.92 |
| **g__Lawsonia** | 0.024975 | 0.12487 | 73961 | 18559 | -4.44 |
| **g__Parabacteroides** | 0.054664 | 0.17562 | 27820 | 129270 | 4.71 |
| **g__Eubacterium** | 0.058539 | 0.17562 | 0 | 24853 | 4.09 |
| **g__Anaeroplasma** | 0.22151 | 0.55377 | 21588 | 14402 | -3.56 |
| **g__Staphylococcus** | 0.2615 | 0.56035 | 3160.1 | 15863 | 3.8 |
| **g__** | 0.42334 | 0.75989 | 5980800 | 5348700 | -5.5 |
| **g__Coprobacillus** | 0.47508 | 0.75989 | 917.71 | 2938 | 3 |
| **g__Ruminococcus** | 0.50659 | 0.75989 | 130710 | 389540 | 5.11 |
| **g__Prevotella** | 0.63095 | 0.86039 | 381100 | 360220 | -4.02 |
| **V1** | 0.74877 | 0.86397 | 150270 | 123990 | -4.12 |
| **g__Lactobacillus** | 0.74877 | 0.86397 | 1812500 | 2023900 | 5.02 |
| **g__Streptococcus** | 0.93607 | 1 | 5503.3 | 2705.6 | -3.15 |
| **g__Bacteroides** | 1 | 1 | 1291800 | 1303900 | 3.78 |
